# The impact of housing conditions on porcine mesenchymal stromal/stem cell populations differ between adipose tissue and skeletal muscle

**DOI:** 10.1101/2021.06.08.447546

**Authors:** Audrey Quéméner, Frédéric Dessauge, Marie-Hélène Perruchot, Nathalie Le Floc’h, Isabelle Louveau

## Abstract

**Background:** In pigs, the ratio between lean mass and fat mass in the carcass determines production efficiency and is strongly influenced by the number and size of cells in tissues. During growth, the increase in the number of cells results from the recruitment of different populations of multipotent mesenchymal stromal/stem cells (MSCs) residing in the tissues. We hypothesized that the impact of hygiene of housing conditions during growth on the proportions of MSCs in adipose tissue and skeletal muscle may differ between pigs with different residual feed intake (RFI), a measure of feed efficiency.

**Methods:** At the age of 12 weeks, Large White pigs from two lines divergently selected for low and high RFI were housed in two contrasting hygiene conditions (good vs poor). After six weeks, pigs were slaughtered (n = 30; 5-9/group). Samples of subcutaneous adipose tissue and longissimus skeletal muscle were collected, and cells from the stromal vascular fraction (SVF), which includes mesenchymal stromal/stem cell populations, were isolated from each tissue. Adipose and muscle cell populations from the SVF were phenotyped by flow cytometry using antibodies that targeted different cell surface markers (CD45 to separate hematopoietic cells from MSCs; CD34, CD38, CD56 and CD140a to identify MSC populations with adipogenic and/or myogenic potential).

**Results:** Adipose tissue and muscle shared some common MSC populations although MSC diversity was higher in muscle than in adipose tissue. In muscle, the CD45^-^CD56^+^CD34^-^CD140a^+^ and CD45^-^CD56^+^CD34^+^CD140a^+^ cell populations were abundant. Of these two cell populations, only the proportions of CD45^-^CD56^+^CD34^+^CD140a^+^ cells increased (P < 0.05) in pigs housed in poor hygiene as compared with pigs in good hygiene conditions. For the CD45^-^CD56^-^CD34^-^ cell population, present in low proportion, there was an interaction between hygiene condition and genetic line (P < 0.05) with a decrease in low RFI pigs housed in poor hygiene conditions. In adipose tissue, the two abundant MSC populations were CD45^-^CD56^-^CD34^-^ and CD45^-^CD56^+^CD34^-^. The proportion of CD45^-^CD56^-^CD34^-^ cells increased (P < 0.05) whereas the proportion of CD45^-^CD56^+^CD34^-^ tended to decrease (P < 0.1) in pigs housed in poor conditions. This study shows that the proportions of some MSC populations were affected by hygiene of housing conditions in a tissue-dependent manner in pigs of both RFI lines. It suggests that MSCs may play a significant role in adipose tissue and skeletal muscle homeostasis and may influence later growth and body composition in growing animals.

## Introduction

During postnatal growth and development, farm animals like pigs are exposed to a diversity of environmental stimuli (Colditz & Hine, 2016). Stressful situations like transport and housing in poor hygiene conditions result in immune system hyper-activation (Stavrakakis et al., 2019). The animal ability to cope with these different challenges will influence the growth of tissues associated with meat production and subsequently their growth efficiency. Fat animals are generally less efficient in the conversion of feed into meat (Sillence, 2004). Moreover, as excess body fat triggers the production of immune and inflammatory cells, the lean-to-fat ratio in the body may influence the ability to fight against infection (Patience, Rossoni-Serão & Gutiérrez, 2015). Hence, a better control of body composition in growing pigs requires a better understanding of the dynamics and flexibility of muscle and adipose tissue growth when the surrounding environment of pigs is modified. The mass of skeletal muscle and adipose tissue is largely determined by the number and size of muscle fibers and adipocytes (Lefaucheur, 2010; Louveau et al., 2016). This increase in tissue mass during pre- and post-natal life results from recruitment of multipotent mesenchymal stromal/stem cells (MSCs) resident in tissues (Dominici et al., 2006). These cells comprise heterogeneous populations of undifferentiated cells able of self-renewal with the capacity to differentiate into cells of the mesenchymal lineage (adipocytes, myocytes or osteocytes for instance). These MSCs have now emerged as relevant targets to control growth and final body composition in livestock (Johnson, 2013; Dodson et al., 2015). It has been shown that the composition of the MSC compartment in adipose tissue is affected by obesity and other disorders in adults (Mihaylova, Sabatini & Yilmaz, 2014; Silva & Baptista, 2019). In growing pigs, we have shown recently that dietary treatments affected the relative proportions of some MSC populations in skeletal muscle and adipose tissue (Perruchot et al., 2020).

The identification of MSC populations is based on the use of a combination of cell surface markers and flow cytometry (Bourin et al., 2013; Relaix et al., 2021). Thus, MSCs are positive for CD105, CD90, CD73 and negative for hematopoietic markers CD11b, CD14, CD34 and CD45. More specifically, both skeletal muscle and adipose tissue cells include resident MSCs that share common cell surface markers. For example, the CD56 marker, also known as a neural cell adhesion molecule (NCAM) and a myogenic marker in muscle (Pisani et al., 2010a; Wilschut et al., 2011; Uezumi et al., 2014), has been also found in MSCs from porcine adipose tissue (Perruchot et al., 2013). Recently, the phenotypic marker CD140a, or platelet derived growth factor receptor alpha (PDGFRα), emerged as an important MSC marker. It has been shown that CD140a^+^ cells derived from muscle or adipose tissue have adipogenic potential (Lee & Granneman, 2012, Uezumi et al., 2010, 2016; Sun, Berry & Olson, 2017). Finally, CD38, one of the main NAD-degrading enzymes in mammalian tissues, has been suggested as a novel marker of committed preadipocytes in mice (Carrière et al., 2017). Other studies have shown that the combination of CD34 and CD56 markers allow the discrimination between myogenic and adipogenic progenitors in human muscle (Pisani et al., 2010a,b). Taken together, these cell surface markers can be used to investigate the potential changes of the relative proportions of MSC populations in muscle and adipose tissue.

Despite the importance of controlling growth and body composition in meat producing animals and the cross-talk between adipose tissue and skeletal muscle throughout life (Crisan et al., 2008; De Carvalho et al., 2019), there is a lack of studies investigating the response of both tissues to different factors. In this context, we propose to characterize and compare MSC populations from adipose and skeletal muscle tissues in response to hygiene of housing conditions in two pig lines divergently selected for Residual Feed Intake (RFI), a measure of feed efficiency (Gilbert et al., 2007). Genetic selection based on production criteria have been suspected to contribute to weaken the immune response and to make pigs more susceptible to behavioral, physiological and immunological problems (Rauw et al., 1998) and infectious diseases (Knap & Doeschl-Wilson, 2020). However, it has been shown that the impact of a challenge based on poor hygiene of housing conditions on growth performance, immune system and health was lower in pigs selected for low RFI (Chatelet et al., 2018). We hypothesized that hygiene of housing conditions will impact the proportions of hematopoietic and/or mesenchymal stromal/stem cells in subcutaneous adipose tissue (SCAT) and skeletal muscle and that the response may differ between pigs with different RFI.

## Materials & Methods

### Ethical approval/declaration

The experiment was performed in the INRAE UE3P experimental facility (Saint-Gilles, France; (doi.org/10.15454/1.5573932732039927E12) in compliance with the ethical standards of the European Community (Directive 2010/63/EU) and was approved by the regional ethical committee (Comité Rennais d’Ethique en matière d’expérimentation animale or CREEA Rennes). The experiment approval number is APAFIS#494–2015082717314985.

### vAnimals

For this experiment, a subset of pigs (n = 30) was selected from a larger study previously described (Chatelet et al., 2018; Sierżant et al., 2019). Growing Large White pigs (males and females) originated from the 8th generation of two lines divergently selected for residual feed intake (RFI), a measure of feed efficiency (Gilbert et al., 2007). Briefly, animals from the high RFI (HRFI) and the low RFI (LRFI) lines were weaned at 4 weeks of age. At 12 weeks, half of the LRFI and HRFI pigs were then assigned to a room with good hygiene conditions and the other half to a room with poor hygiene conditions. Good housing conditions included room cleaning, disinfection, and adoption of strict biosecurity precautions. In contrast, poor hygiene conditions consisted of no cleaning nor sanitation of the room after the previous occupation by non-experimental pigs. All pigs were housed in individual pens (85 × 265 cm) in concrete floor throughout the experimental period and were fed *ad libitum* a standard diet formulated to meet or exceed the nutritional requirement of growing pigs (9.47 MJ of net energy/kg, starch 44.2%; fat 3.1%; crude protein 15.3%. They had free access to water. After six weeks and after an overnight fast, animals (n = 30; 5-9/experimental group) were euthanized by electrical stunning and exsanguination. Immediately after slaughter, samples of subcutaneous dorsal adipose tissue (SCAT) and of *longissimus* muscle were excised and placed in 37°C Krebs-Ringer-bicarbonate-HEPES buffer for cell isolation (Perruchot et al., 2013).

### Adipose cell isolation

Cells were isolated from a sample of fresh dorsal subcutaneous adipose tissue by collagenase digestion as previously described (De clercq et al., 1997; Perruchot et al., 2013). Briefly, fresh adipose tissue was cut into small pieces and dissociated by enzymatic digestion with collagenase II and XI (800 U/mg; Sigma, Saint-Quentin-Fallavier, France) under shaking in a dry bath for 45 min at 37°C. Then, a centrifugation at 400 g for 10 min was performed to separate floating adipocytes from the pellet of stromal vascular fraction (SVF) cells. After resuspension, SVF cells were successively filtered through a 200-µm and 25-µm nylon membranes (Dutscher, Brumath, France). After isolation, SVF cells were placed in 90% Fetal Calf Serum (FCS) and 10% dimethyl sulfoxide (1–2 million cells per mL) and frozen at - 150°C.

### Muscle cell isolation

Fresh *longissimus* muscle tissue devoid of visible connective tissue was finely chopped and digested for 3 × 20 min in a Ca^2+^-free medium containing 0.25% trypsin (Invitrogen, Cergy-Pontoise, France), 1.5 mg/mL collagenase II (PAA Laboratoires, Les Mureaux, France), and 0.1% DNAse (Sigma) in a shaking water bath at 37°C. After centrifugation at 800 g for 10 min at 4°C, cell pellet was re-suspended in ice-cold proliferation growth medium (PGM) containing DMEM supplemented with 10% FCS (Invitrogen), 10% horse serum (Invitrogen), and Penicillin 6,25 UI/ml -Streptomycin 6,25 µg/ml, (Fisher Scientific, Illkirch, France). Next, cell suspension was successively filtered through a 200-µm and 50-µm Nylon mesh (Dutscher). After their collection in ice-cold PGM, isolated cells were placed in 90% Fetal Calf Serum (FCS) and 10% dimethyl sulfoxide (1-2 million cells per mL) and frozen at - 150°C.

### Flow cytometry analysis

Individual vials of cryopreserved cells were thawed in a 37°C water bath (2-3 min). After thawing, cells were suspended in 10 mL of FCS (Sigma). After centrifugation at 500 g for 5 min at room temperature (RT), cells were suspended in 10 mL of DPBS (Fisher Scientific). Cells were then counted with the TC20™ Automated Cell Counter (Bio-Rad, Marnes-la-Coquette, France). Next, cells were centrifuged at 500 g for 5 min at RT and suspended in 10 mL of FACS buffer. This FACS buffer contained MACSQuant Running Buffer (Miltenyi Biotec, Paris, France) and 0.5% of MACS® BSA Stock Solution (Miltenyi Biotec). Cells were dispensed at 50,000 cells for SCAT and 500,000 cells for muscle in each round bottom tube of 5 mL (Sarstedt, Nümbrecht, Germany). After centrifugation at 500 g for 5 min at RT, cells were suspended in 100 µL of FACS buffer. Cells were incubated in the dark on ice for 15 min with labeled monoclonal or polyclonal antibodies coupled to different fluorochromes: allophycocyanine (APC), brilliant violet 421 (BV421), fluorescein isothiocyanate (FITC), phycoerythrin (PE), phycoerythrin-vio770 (PE-Vio770) (Table 1). After incubation with antibodies, cells were washed in FACS buffer and centrifuged at 500 g for 5 min at 4°C. Labeled cells were then suspended in 400 µL of FACS buffer. Labeled cells were analyzed using a MACSQuant Analyzer 10 flow cytometer (Miltenyi Biotec), which was calibrated daily. A minimum of 20,000 events was acquired per sample for SCAT and 200,000 events for muscle. Propidium iodide (Miltenyi Biotec) was used to exclude dead cells. Flow cytometry analyses were performed with appropriate isotype matching negative and Flow Minus One (FMO) controls. Data were analyzed using the FlowLogic software V8 (Inivai, Mentone, Victoria, Australia).

**Table 1:** List of antibodies used for flow cytometry analysis

| Antigen | Antibody | Reference | Dilution |
| --- | --- | --- | --- |
| **CD34** | CD34-PE (clone EP373Y) | Abcam (ab223930) | 1:100 |
| Isotype control | IgG-PE (clone EPR25A) | Abcam (ab209478) | 1:100 |
| **CD38** | CD38-PE-Vio770 (clone REA 671) | Miltenyi Biotec (130-119-159) | 1:25 |
| Isotype control | IgG1-PE-vio770 (REA 293) | Miltenyi Biotec (130-113-440) | 1:25 |
| **CD45** | CD45-FITC (clone K252.1E4) | (BIO RAD) MCA1222F | 1:10 |
| Isotype control | IgG1-FITC (clone MCA928F) | (BIO RAD) MCA928F | 1:10 |
| **CD56** | CD56-BV421 (clone NCAM16.2) | BD Biosciences (562751) | 1:10 |
| Isotype control | (BALB/c) IgG2b, κ | BD Biosciences (562748) | 1:10 |
| **CD140a** | CD140a-APC (clone REA 637) | Miltenyi Biotec (130-122-942) | 1:10 |
| Isotype control | IgG1-APC (REA 293) | Miltenyi Biotec (130-113-434) | 1:50 |

### Statistical analyses

Statistical analyses were performed using the GraphPad Prism software (version 8.4.3 (686), San Diego, CA, USA). Residual plots for parameters significantly different between groups can be found as Supplementary Figure S1. The data were subjected to the two-way analysis of variance (ANOVA). Hygiene of housing conditions, RFI line and their interactions were considered as fixed effects. Results are presented as mean ± standard error of mean (SEM). Differences were considered statistically significant at P ≤ 0.05 and were discussed as a trend for 0.05 < P < 0.1.

## Results

### Growth performance and body composition of the studied groups

Data for the subset of pigs investigated in the current study are shown in Table 2. There was no significant interaction between housing conditions and RFI lines for pig performance and their body composition. Final body weights, average daily gain and backfat weight did not differ between the experimental groups (P > 0.1). Interestingly, irrespective of housing conditions, the relative loin weight was lower in HRFI pigs than in LRFI pigs (P = 0.005). Finally, irrespective of the RFI line, the loin/backfat ratio was higher (P = 0.045) in pigs raised in poor conditions than in pigs raised in good conditions.

**Table 2:** Growth performance of low (LRFI) and high (HRFI) residual feed intake pigs housed in good or poor hygiene conditions for six weeks

|  | Good |  | Poor |  | *P-values*<sup>1</sup> |  |  |
| --- | --- | --- | --- | --- | --- | --- | --- |
|  | LRFI | HRFI | LRFI | HRFI | Hyg | Line | Hyg x Line |
| N | 8 | 7 | 9 | 6 |  |  |  |
| Initial body weight (kg) | 27.8 ± 0.7 | 25.7 ± 1.7 | 28.2 ± 0.7 | 25 ± 1.4 | 0.402 | 0.495 | 0.289 |
| Final body weight (kg) | 54.6 ± 2.6 | 51.1 ± 1.9 | 54.0 ± 3.4 | 49.8 ± 3.3 | 0.783 | 0.174 | 0.958 |
| Average daily gain (g/day) | 626 ± 67 | 590 ± 59 | 603 ± 84 | 564 ± 48 | 0.750 | 0.614 | 0.919 |
| Relative loin weight (%) | 27.6 ± 0.3 | 27.0 ± 0.3 | 28.4 ± 0.2 | 27.2 ± 0.5 | 0.137 | **0.005** | 0.274 |
| Relative backfat weight (%) | 4.9 ± 0.2 | 5.0 ± 0.1 | 4.6 ± 0.3 | 4.6 ± 0.3 | 0.125 | 0.903 | 0.962 |
| Loin/backfat ratio | 5.6 ± 0.2 | 5.5 ± 0.2 | 6.5 ± 0.5 | 6.0 ± 0.3 | **0.045** | 0.322 | 0.638 |

Values are means ± SEM. Boldface highlights significant differences ( $P \leq 0.05$ ).
<sup>1</sup>Probability values for the effect of hygiene conditions (Hyg), genetic lines (Line), and their interaction.

### Description of the phenotyping strategy

To identify the cell populations specific and common to SCAT and muscle, we focused our analysis on a panel of five cell surface markers: CD45, CD34, CD38, CD56 and CD140a (Figure 1).

**Figure 1:**
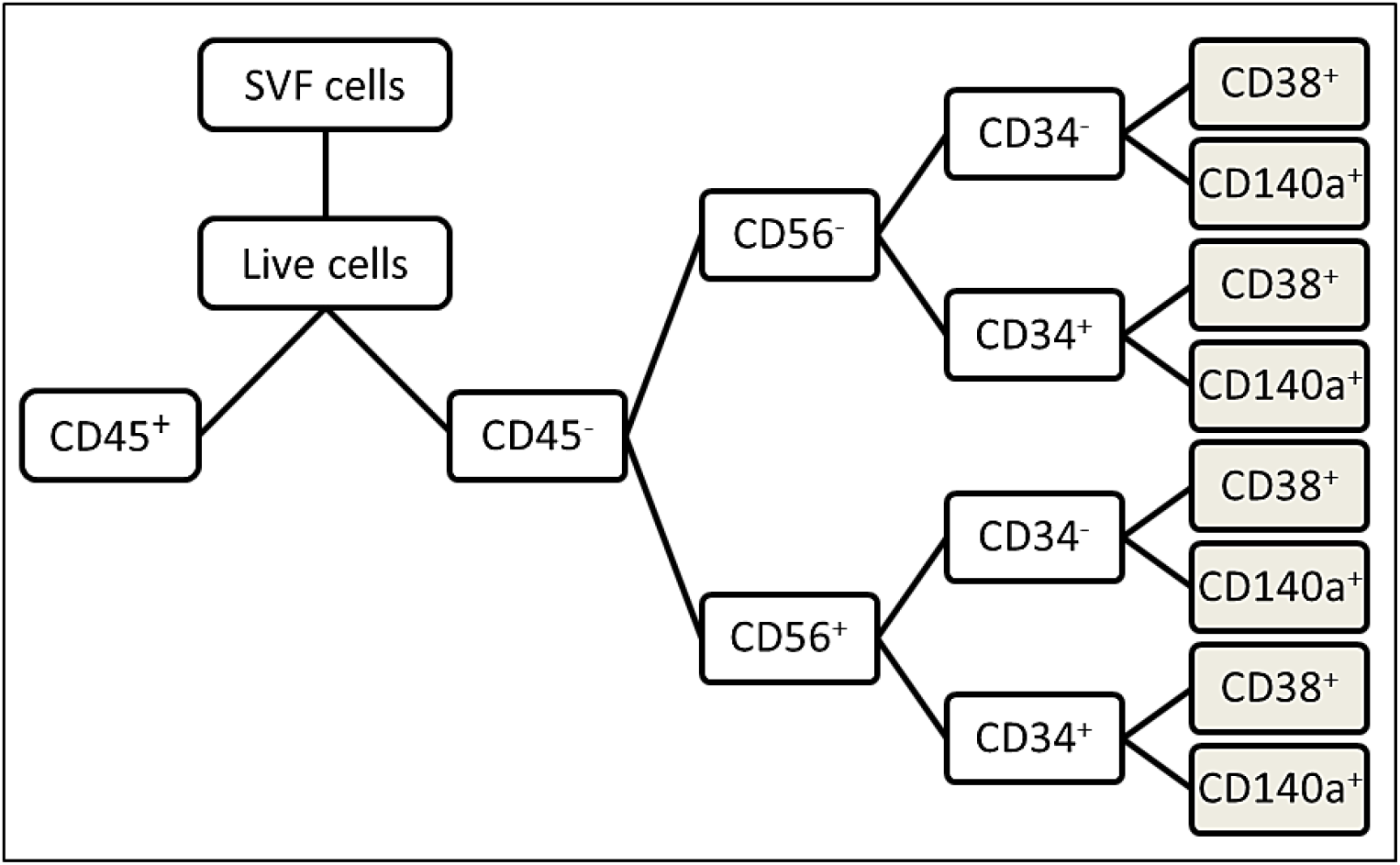
Diagram of the strategy used for cell immunophenotyping by flow cytometry. SVF cells were isolated from adipose tissue and skeletal muscle of growing pigs with low and high residual feed intake and housed in good or poor hygiene conditions for six weeks. A common analytical strategy was used for both tissues. First, live cells were selected from the total SVF cell pool. Then, a first discrimination based on CD45 marker was carried out in order to separate hematopoietic cells (CD45+) from MSCs (CD45-).

Then, a phenotyping at the level of four lineage was performed on the MSCs with the CD56, CD34 and CD38 or CD140a (white and brown squares) markers. Adipose tissue was only phenotyped at the level of triple lineage because of its low quantities of CD45-CD56-CD34+ and CD45-CD56+CD34+ cells (white squares).

First, gating was performed on live cells that were negative for the CD45 in order to remove hematopoietic cells positive for the CD45 marker. Next, we analyzed the following combinations in SCAT and muscle: CD45^-^CD34^+^, CD45^-^CD38^+^, CD45^-^CD56^-^, CD45^-^CD56^+^, CD45^-^CD140a^+^. Then, we performed a deeper phenotyping for cell populations predominant in both SCAT and muscle, respectively CD45^-^CD56^-^ and CD45^-^CD56^+^ (Figures 1, 2 and 3). Based on the literature, we evaluated within these two cell populations, the expression of the CD34 marker as named: CD45^-^CD56^-^CD34^-^, CD45^-^CD56^-^CD34^+^, CD45^-^ CD56^+^CD34^-^ and CD45^-^CD56^+^CD34^+^ populations. Given the low proportion (<0.5%) of some cell populations in SCAT, we only analyzed the expression of the CD38 and CD140a cell surface markers in muscle in order to highlight the proportion of the following populations: CD45^-^CD56^-^CD34^-^CD38^+^, CD45^-^CD56^-^CD34^-^ CD140a^+^, CD45^-^CD56^-^CD34^+^CD38^+^, CD45^-^CD56^-^CD34^+^CD140a ^+^, CD45^-^CD56^+^CD34^-^CD38^+^, CD45^-^CD56^+^CD34^-^ CD140a^+^, CD45^-^CD56^+^CD34^+^CD38^+^ and CD45^-^CD56^+^CD34^+^CD140a^+^. Altogether, this gating strategy allowed us to highlight specific and common cell populations in both SCAT and muscle.

**Figure 2:**
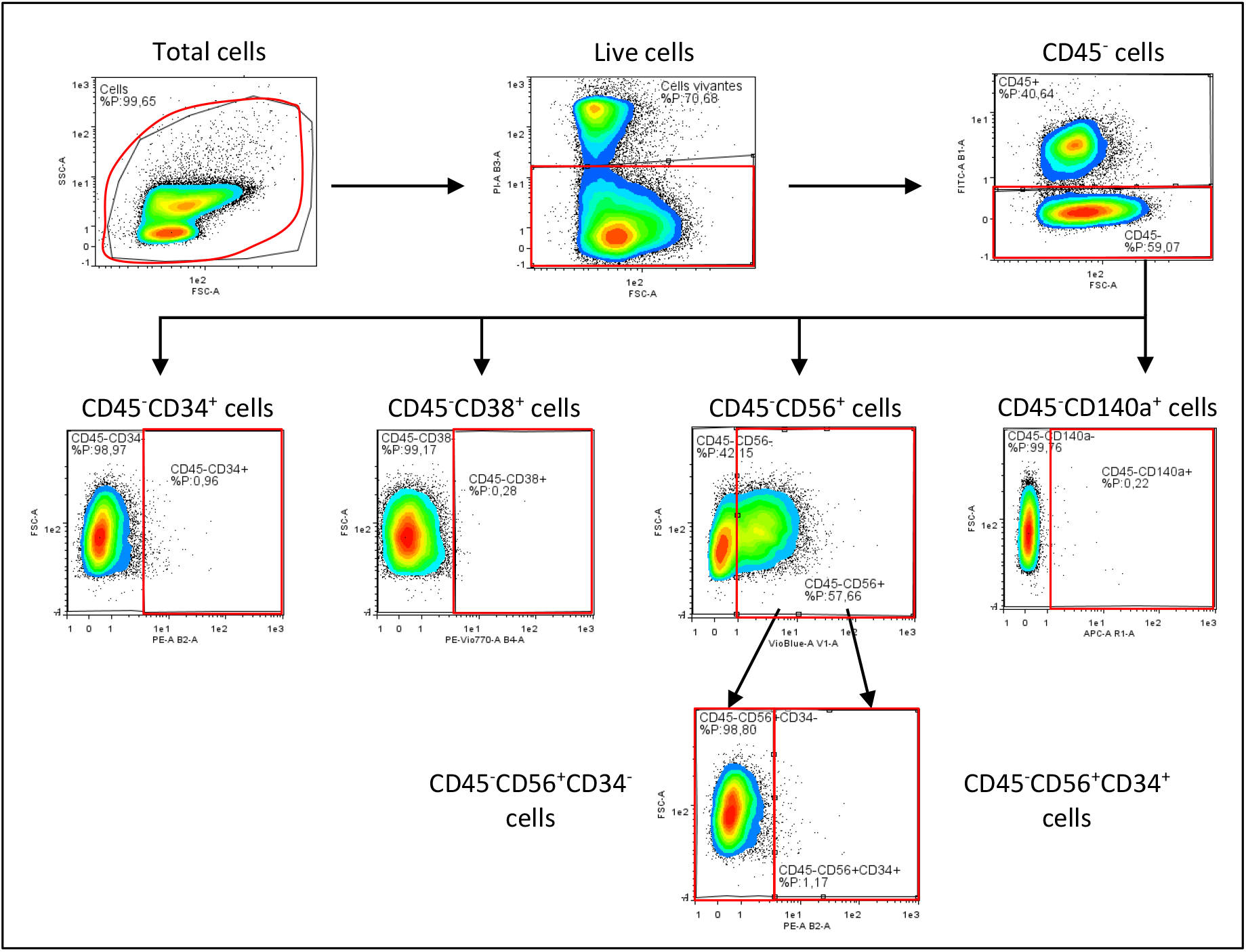
Representative flow cytometry plots illustrating the immunophenotyping strategy used for mesenchymal stromal/stem cells isolated from adipose subcutaneous tissue of growing pigs. The captions correspond to the cell populations highlighted in red. In addition, the red boxes delimit the positivity threshold of studied cell populations established according to the appropriate isotype and Fluorescence Minus One controls.

**Figure 3:**
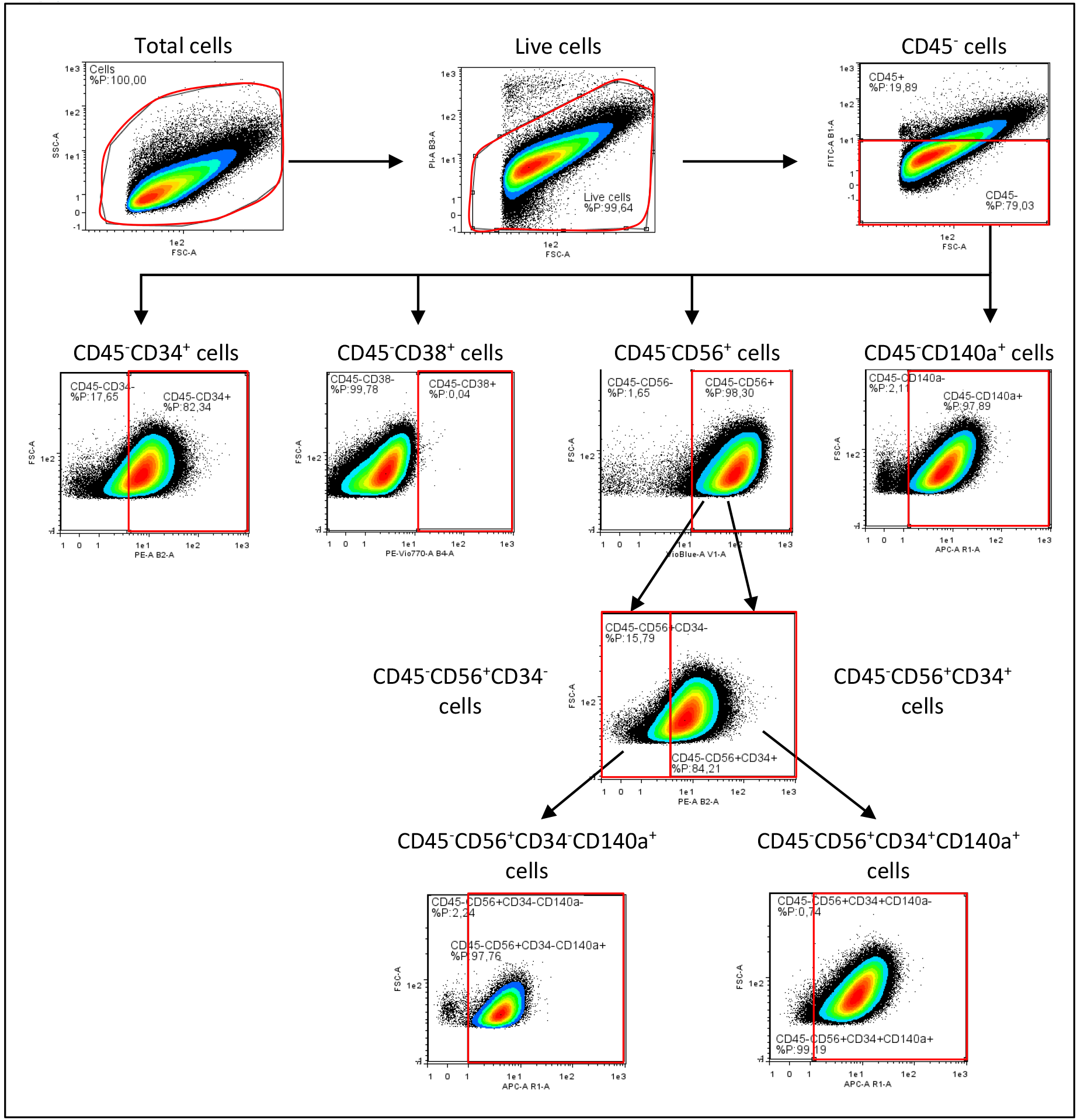
Representative flow cytometry plots illustrating the immunophenotyping strategy used for mesenchymal stromal/stem cells isolated from *longissimus* muscle tissue of growing pigs. The captions correspond to the cell populations highlighted in red. In addition, the red boxes delimit the positivity threshold of studied cell populations established according to the appropriate isotype and Fluorescence Minus One controls.

### Analysis of cell populations identified in subcutaneous adipose tissue

There was a substantial proportion of CD45^+^ cells (23.1%) in SCAT with no difference between the four experimental groups (Table 3). We next analyzed the occurrence of co-staining for the four cell surface markers CD56, CD34, CD38 and CD140a in order to further delineate the different populations within CD45^-^ cells dissociated from SCAT (Figures 1 and 2). With triple staining, the proportions of the CD45^-^CD56^-^CD34^-^ population was higher (+11.7%, P = 0.034) in poor than in good hygiene of housing conditions whatever the RFI line (Figure 4A) whereas the proportion of the CD45^-^CD56^+^CD34^-^ population tended to decrease (−17.2%, P = 0.099) in poor hygiene conditions compared with good conditions (Figure 4C). Of note, the other cell populations with double or triple staining, such as CD45^-^CD56^+^CD34^+^ cells (Figure 4E) were present in very low proportions (<1%) in SCAT (Table 3).

**Table 3:** Percentage of cell populations in subcutaneous adipose tissue of low (LRFI) and high (HRFI) residual feed intake pigs housed in good or poor hygiene conditions for six weeks

| Cell populations | Good |  | Poor |  | *P-values*<sup>1</sup> |  |  |
| --- | --- | --- | --- | --- | --- | --- | --- |
|  | LRFI | HRFI | LRFI | HRFI | Hyg | Line | Hyg x Line |
| **N** | 7 | 7 | 9 | 6 |  |  |  |
| **CD45<sup>+</sup>** | 18.2 ± 7.1 | 22.7 ± 7.1 | 26.7 ± 5.1 | 24.1 ± 5.4 | 0.434 | 0.880 | 0.571 |
| **CD45<sup>-</sup>** | 81.5 ± 7.1 | 77.0 ± 7.1 | 72.7 ± 5.2 | 75.7 ± 5.4 | 0.429 | 0.905 | 0.558 |
| **CD45<sup>-</sup>CD34<sup>+</sup>** | 0.5 ± 0.2 | 0.3 ± 0.1 | 0.4 ± 0.1 | 0.6 ± 0.1 | 0.530 | 0.844 | *0.071* |
| **CD45<sup>-</sup>CD38<sup>+</sup>** | 0.3 ± 0.1 | 0.2 ± 0.1 | 0.4 ± 0.1 | 0.5 ± 0.1 | *0.082* | 0.779 | 0.212 |
| **CD45<sup>-</sup>CD140a<sup>+</sup>** | 0.5 ± 0.1 | 0.1 ± 0.02 | 0.3 ± 0.1 | 0.3 ± 0.1 | 0.607 | *0.059* | **0.036** |
| **CD45<sup>-</sup>CD56<sup>-</sup>** | 27.4 ± 6.0 | 29.5 ± 5.2 | 42.4 ± 3.7 | 36.8 ± 5.1 | **0.035** | 0.724 | 0.448 |
| **CD45<sup>-</sup>CD56<sup>-</sup>CD34<sup>-</sup>** | 26.8 ± 6.0 | 29.2 ± 5.2 | 41.9 ± 3.7 | 36.4 ± 5.0 | **0.034** | 0.757 | 0.442 |
| **CD45<sup>-</sup>CD56<sup>-</sup>CD34<sup>+</sup>** | 0.6 ± 0.2 | 0.2 ± 0.04 | 0.5 ± 0.2 | 0.3 ± 0.1 | 0.746 | 0.106 | 0.591 |
| **CD45<sup>-</sup>CD56<sup>+</sup>** | 53.9 ± 12.2 | 47.3 ± 9.9 | 30.1 ± 6.2 | 38.6 ± 10.0 | 0.101 | 0.920 | 0.437 |
| **CD45<sup>-</sup>CD56<sup>+</sup>CD34<sup>-</sup>** | 53.7 ± 12.3 | 47.1 ± 10.0 | 29.8 ± 6.2 | 38.3 ± 10.0 | *0.099* | 0.922 | 0.439 |
| **CD45<sup>-</sup>CD56<sup>+</sup>CD34<sup>+</sup>** | 0.2 ± 0.1 | 0.2 ± 0.1 | 0.2 ± 0.1 | 0.3 ± 0.1 | 0.230 | 0.653 | 0.693 |

Values are means ± SEM. Boldface highlights significant differences ( $P \leq 0.05$ ) and italicized characters show a trend ( $0.05 < P < 0.10$ ).
<sup>1</sup>Probability values for the effect of hygiene conditions (Hyg), genetic lines (Line), and their interaction.

**Figure 4:**
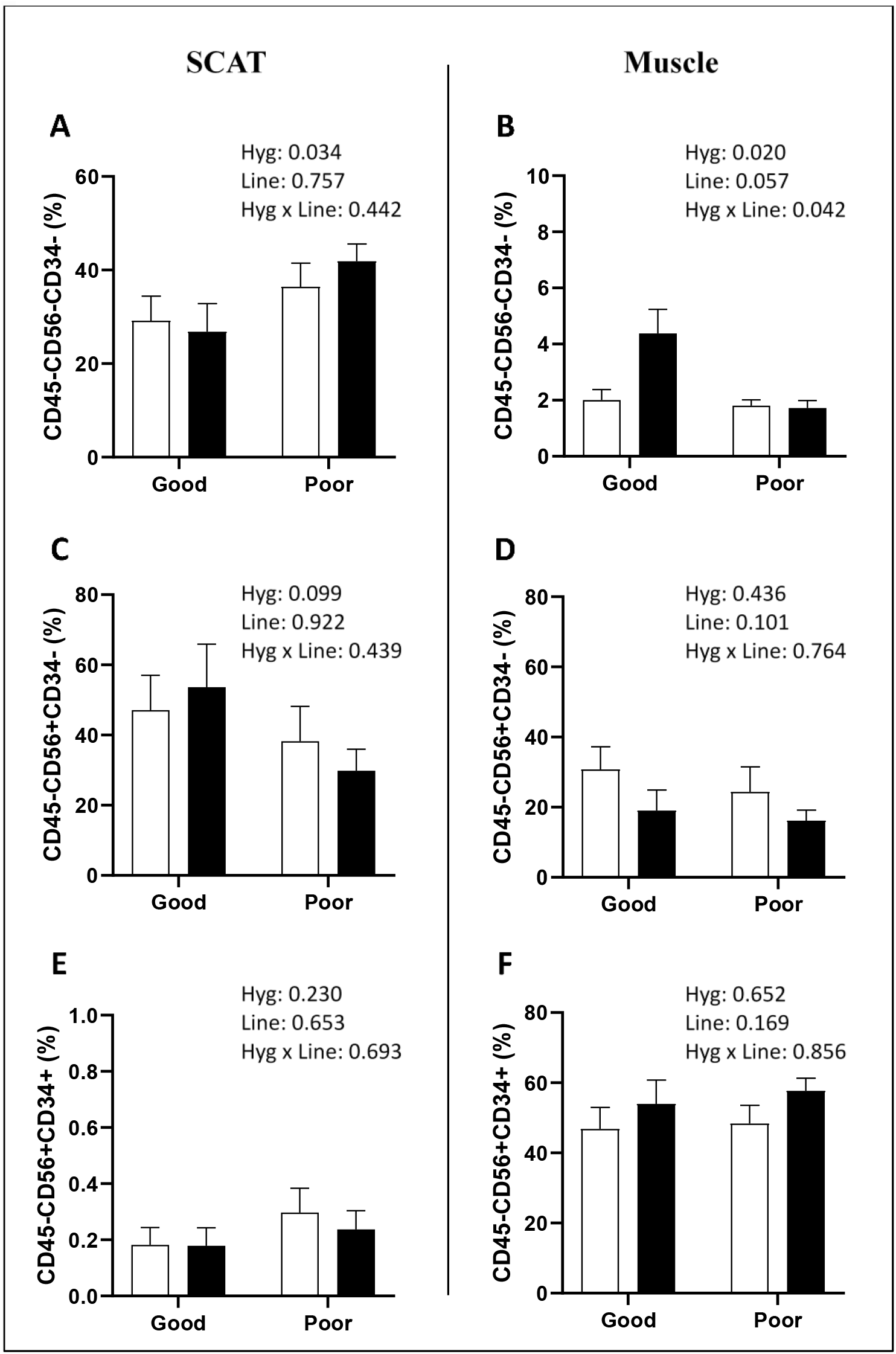
Percentage of mesenchymal stromal/stem cell populations in subcutaneous adipose tissue (SCAT) and *longissimus* muscle from low (LRFI, black bar) and high (HRFI, white bar) residual feed intake pigs housed in good or poor hygiene conditions. Percentage of cell populations are expressed as means ± SEM (n= 5-9/experimental group).

### Analysis of cell populations identified in skeletal muscle

The proportion of CD45^+^ cells in *longissimus* muscle was in the range of 20-23% (Table 4) and increased (P = 0.020) in pigs housed in poor conditions compared with pigs housed in good conditions for both RFI lines. Next, we analyzed the expression of CD34, CD38, CD140a and CD56 within the CD45^-^ cell population (Figures 1 and 3). Despite their high proportions (>50%) in muscle, the CD45^-^CD34^+^ and CD45^-^CD140a^+^ cell populations did not differ significantly between the four groups. Furthermore, there was an interaction for the CD45^-^CD56^-^ cell population (P = 0.011). Interestingly, within this CD45^-^CD56^-^ cell population, we analyzed the expressions of CD34 and CD140a. We also observed a significant interaction for the CD45^-^CD56^-^CD34^-^ cell population (P = 0.042) (Figure 4B). For the CD45^-^CD56^-^CD34^-^CD140a^+^ cells, their proportion tended to decrease (P = 0.072) in poor compared with good hygiene conditions. Moreover, within the CD45^-^CD56^+^ cell population, the proportion of CD45^-^CD56^+^CD34^-^ cells tended to be higher in HRFI pigs (+10%, P = 0.101) than in LRFI pigs (Figure 4D). Concerning the CD45^-^CD56^+^CD34^+^ cells, their proportions do not vary whatever the genetic line or the housing conditions (Figure 4F). Finally, by adding the CD140a marker in the analysis, we highlighted that, irrespective of the genetic line, the CD45^-^CD56^+^CD34^+^CD140a^+^ cells increased (+ 11.3%) in poor conditions (P = 0.052) compared with good conditions (Table 4).

**Table 4:**
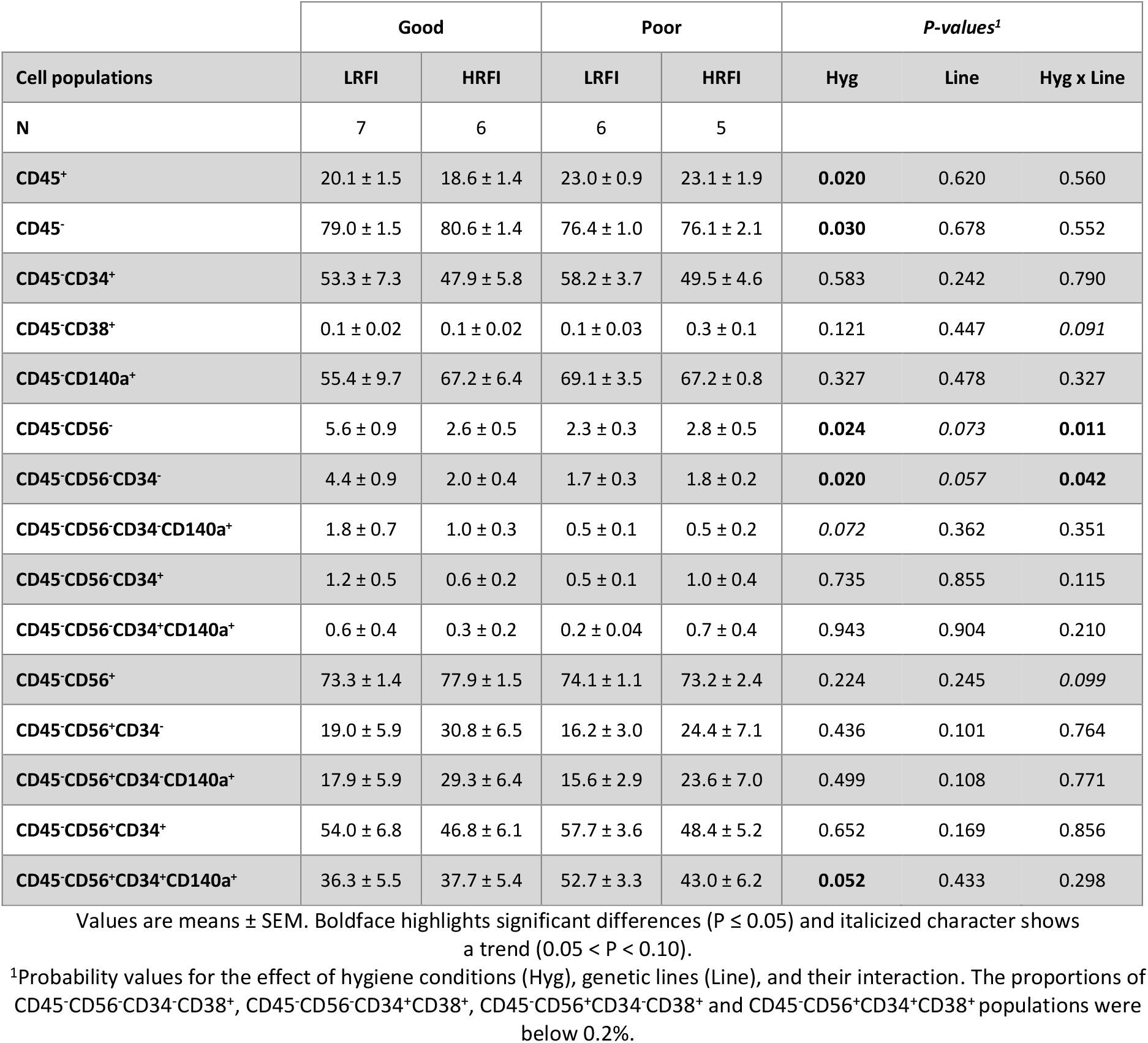
Percentage of cell populations in *longissimus* muscle of low (LRFI) and high (HRFI) residual feed intake pigs housed in good or poor hygiene conditions for six weeks

## Discussion

The current study was conducted to further enrich our knowledge on mesenchymal stromal/stem cell populations resident in muscle and adipose tissue, two tissues of importance for body composition and meat quality traits in pigs. We determined whether the exposure of growing HRFI and LRFI pigs to poor hygiene of housing conditions impact the proportions of these cells. It further documents the phenotype of MSCs. Whatever the species considered, the characterization of MSCs in muscle and adipose tissue is fairly recent and is conventionally carried out using flow cytometry which is based on the use of antibodies directed against cell surface proteins (Bourin et al., 2013; Relaix et al., 2021). For the current cell immunophenotyping, we have used five well-known cell surface markers (CD45, CD56, CD34, CD38 and CD140a) to identify specific and common CD45^-^ MSC populations in both tissues. Moreover, we determined the proportions of hematopoietic cells in both tissues (Han et al., 2021), the CD45^+^ cells, a population known to be increased after injury in muscle (Wosczyna et al., 2019). Our major finding is that the proportions of several cell populations were affected by hygiene of housing conditions in a tissue-dependent manner in pigs of both RFI lines.

Even though health and growth performance of the most feed efficient LRFI pigs was less impaired by poor hygiene conditions (Chatelet et al., 2018), our findings at the cellular levels do not support a significant relationship between MSCs and the better coping ability of LRFI compared with HRFI pigs. In the current study, there was only a significant interaction for the CD45^-^CD56^-^CD34^-^ population in muscle, a relatively small population. For the other cell populations, there was no significant difference between the two lines. This differs slightly from a previous study showing some differences in some cell populations between the two RFI lines (Perruchot et al., 2020). In this latter study, castrated males with higher body weights were investigated instead of males and females in the present study. These differences may contribute to the difference between the two studies.

The current study clearly shows the presence of a relative high proportion of CD45^+^ cells suggesting the presence of hematopoietic cells in both muscle and adipose tissue as shown in previous studies (Cousin et al., 2016; Wosczyna et al., 2019). Our study also indicates that the proportion of CD45^+^ cells in muscle increased with the exposure of pigs to poor hygiene of housing conditions. This increase may be associated with the inflammation induced by the poor hygiene of housing conditions. The stimulation of the immune system of pigs housed in these conditions was assessed by an increase in plasma haptoglobin concentrations, an acute phase protein (APP), and in white blood cell and granulocyte counts three weeks after the beginning of the challenge (Chatelet et al., 2018). The origin of this increase in CD45^+^ cells remains unknown. It could result from an increase in CD45^+^ resident cells in skeletal muscle. Alternatively, it could come from other tissues as reported for MSCs in a recent review (Girousse et al., 2021). A study based on the investigation of MSCs from adipose tissue and peripheral blood of Iberian pigs supports an *in vitro* migration ability of these cells (Calle et al., 2018). Resident MSCs could circulate toward distant inflamed tissues to allow their repair/regeneration. Hence, the observed increase in CD45^+^ cells may be associated with changes in local secretory environment and may affect properties of other muscle cells and affect subsequent growth.

As described previously (Bourin et al., 2013; Girousse et al., 2021; Relaix et al., 2021), there was a diversity of MSC populations in skeletal muscle and adipose tissue of the different groups of animals. We confirmed that CD45^-^CD56^+^ cells are predominant in both muscle and adipose tissue as observed before (Perruchot et al., 2013). In porcine muscle, these cells are mainly myogenic (Perruchot et al., 2013). In human and pig, CD45^-^CD56^+^CD34^-^ were identified as satellite cells (Wilschut et al., 2008; Pisani et al., 2010b; Lewis et al., 2014). In adipose tissue, the presence of CD45^-^CD56^+^ is much less documented. We further highlighted a higher cell diversity in muscle than in adipose tissue. There was especially a high proportion of CD34^+^ and CD140a^+^ cells within the CD45^-^ cells. In pigs, CD45^-^CD34^+^ cells have been previously identified in muscle as PW1^+^ interstitial cells (PICs) with an ability to differentiate into skeletal myoblast/myotubes, smooth muscle, and endothelial cells (Lewis et al., 2014). The CD45^-^CD140a^+^ cells were also located in the interstitial space of muscle. In normal skeletal muscle and damaged skeletal muscle, CD140a^+^ cells exhibit fibrogenic and adipogenic potentials and were named fibro/adipogenic progenitors (FAPs; Uezumi et al., 2010, 2014; Uezumi, Ikemoto-Uezumi & Tsuchida, 2014). In muscle of growing pigs, we have shown that CD56^+^CD140a^+^ and CD56^+^CD140a^-^ were mainly myogenic (Perruchot et al., 2020). Muscle stem cell heterogeneity has been described at several other levels, including transcription factor, clonal capacity, metabolism, age dependent myogenic potential, and recently functional response to environmental stress (Cho & Doles, 2019; Relaix et al., 2021). This multitude of resident stem cells contributes likely together to the remarkable regeneration capacities of muscle after injury (Mackey et al., 2017; Relaix et al., 2021). We also included the CD38 marker in our analysis to determine whether it could be used to identify committed preadipocytes as described in mice (Carrière et al., 2017). However, the very low number of cells expressing this marker does not support these findings. The discrepancy between the two studies may be related to the fact that the CD38 marker was detected in cells from abdominal adipose tissue, an adipose depot exhibiting a greater expression of CD38 than in SCAT.

We highlighted for the first time that the environment influence MSCs in growing pigs. The CD45^-^CD56^+^CD34^+^CD140a^+^ cell population, one of the abundant populations in muscle, increased clearly after an exposure of pigs to poor hygiene of housing conditions. We previously found that CD56^+^CD140a^+^cells were myogenic (Perruchot et al., 2020). With the use of the CD34 and CD56 markers, CD34^+^ cells derived from human muscle were described to be myo-adipogenic bipotent progenitors (Pisani et al., 2010a,b). Thus, we can hypothesize that these cells exhibit myo/adipogenic potentials. It is interesting to note that CD56^+^CD140a^+^ cells were also affected by a high-fat/high-fiber diet in porcine muscle (Perruchot et al., 2020). This suggests that these cells may respond to changes of different factors. The current study further indicates that there was an interaction between hygiene conditions and RFI lines for CD45^-^CD56^-^ cells including CD45^-^CD56^-^CD34^-^ cells representing a low proportion of cells in muscle. There was a decrease of cells in poor hygiene condition only in LRFI pigs. These cells may be perivascular cells as described for CD146^+^CD34^−^CD45^−^CD56^−^ cells in multiple human tissues including muscle (Crisan et al., 2008; Shenoy & Bose, 2018). These cells may have a myo-adipogenic potential. Indeed, we have shown that CD56^-^CD140a^+^ cells in porcine muscle exhibited both adipogenic and myogenic potentials (Perruchot et al., 2020). Moreover, Shenoy & Bose (2018) showed that pericytes from mouse liver were able to regenerate muscle fibers in muscle of dystrophic (Mdx) mice. In addition, it has been shown in skeletal muscle of obese humans that the CD56^-^ cell fraction gives rise to white adipocytes (Laurens et al., 2016). Other studies have reported the existence of adipogenic progenitors, not expressing the CD56 marker, in skeletal muscle MSCs derived from lean individuals in humans (Vauchez et al., 2009; Lecourt et al., 2010). Altogether, available data obtained in pigs and humans are consistent with a myo-adipogenic potential of the CD45^-^CD56^-^CD34^-^ cells.

With the combination of cell surface markers used in the current study, adipose tissue cell composition is less complex than skeletal muscle with two abundant CD45^-^ cell populations, the CD45^-^CD56^-^CD34^-^ and the CD45^-^CD56^+^CD34^-^ populations. The proportion of CD45^-^CD56^-^CD34^-^ cells increased whereas the proportion of CD45^-^CD56^+^CD34^-^ tended to decrease in degraded hygiene conditions. The presence of CD45^-^CD56^+^CD34^-^ cells in adipose tissue is poorly documented in the literature. In two previous studies, we provided evidence that CD56^+^ cells from subcutaneous adipose tissue have an adipogenic potential (Perruchot et al., 2013, 2020). Some groups did not find any CD56^+^ cells in adipose tissue, while others attribute the expression of CD56 to natural killer cells (CD45^+^). Overall, there is no consensus on the expression of the CD56 marker in adipose tissue within the CD45^-^ cells. Moreover, the increase in CD45^-^CD56^-^CD34^-^ cells in poor hygiene condition may increase later adipose tissue growth in this group of pigs. Another possible hypothesis would be that the poor hygiene condition of pig housing may lead to a delay in the development of adipose tissue, and therefore the pool of CD45^-^CD56^-^CD34^-^ cells, potential adipocyte precursors, would be less used in these animals compared with those housed in good condition. Altogether, these findings are consistent with studies demonstrating that the composition of the MSC compartment of adipose tissue and skeletal muscle can be affected by several factors including nutritional factors (Perruchot et al., 2020) during growth.

## Conclusions

The findings gathered in this study clearly show that the relative proportions of hematopoietic and of some mesenchymal stromal/stem cell populations were affected by hygiene of housing conditions in a tissue-dependent manner in pigs of both RFI lines. It further indicates that these cell populations may play a significant role in adipose tissue and skeletal muscle homeostasis and may influence later growth and body composition in growing animals. Thus, further investigations are needed to determine the functions of the different cell populations and to get a better understanding of cell interactions in both tissues.

## Data accessibility

Data are available online: https://doi.org/10.15454/IXXVBB

## Supplementary material

Supplementary Figure S1 is available online: https://doi.org/10.15454/IXXVBB

## Acknowledgements

The authors are very grateful to the staff of the experimental pig facilities from INRAE at UE3P (Saint-Gilles, France) for animal care and slaughtering. They are also grateful to Hélène Gilbert (INRAE, UMR GenePhySe, Toulouse, France) for selection of the pig RFI lines. The authors also wish to thank Frédérique Mayeur and Christine Tréfeu (PEGASE, Saint-Gilles, France) for their contribution to the isolation of adipose and muscle cells. Version 3 of this preprint has been peer-reviewed and recommended by Peer Community In Animal Science (https://doi.org/10.24072/pci.animsci.100011).

